# Stochasticity in multi-phosphorylation and quasi steady state approximation in stochastic simulation

**DOI:** 10.1101/392951

**Authors:** S. Das, D. Barik

## Abstract

Quantitative and qualitative nature of chemical noise propagation in a network of chemical reactions depend crucially on the topology of reaction networks. Multisite reversible phosphorylation-dephosphorylation of target proteins is one such recurrently found topology in various cellular networks regulating key functions in living cells. Here we analytically calculated the stochasticity in multistep reversible chemical reactions by determining variance of phosphorylated species at the steady state using linear noise approximation. We investigated the dependence of variance on the rate parameters in the reaction chain and the number of phosphorylation sites on the species. Assuming a quasi steady state approximation on the multistep reactions, originating from the disparity in time scales in the network, we propose a simulation scheme for coupled chemical reactions to improve the computational efficiency of stochastic simulation of the network. We performed case studies on signal transduction cascade and positive feedback loop with bistability to show the accuracy and efficiency of the method.

## Introduction

An isogenic population of cells, both from single and multi-cellular organisms, in an identical environmental condition show remarkable cell-to-cell variability in protein abundance, cell cycle properties, cell size and shape, timescales of key signaling events resulting in population heterogeneity (1-7). The population heterogeneity is due to the inevitable intrinsic and extrinsic noise, collectively known as chemical noise, affecting chemical reactions inside living cells(8). While chemical noise may act as nuisance, it has potential beneficial roles in cellular physiology and functions (9-11). While intrinsic noise originates from the low molecular abundances of chemical species and produces stochastic trajectories of products over time, extrinsic noise originates from global factors such as variation in cell volume or size, cell cycle phases, abundances of regulatory molecules such as transcription factors affecting all chemical reactions in a given cell equally (12). Using probabilistic description of chemical reactions, theoretical models of gene expression noise were able to quantitatively explain many experimental observations on protein noise (13-15). However cellular functions are regulated by chemical reaction networks involving many proteins and these networks often consists of small regulatory network motifs with distinct properties (16). Therefore further investigations were carried out to understand the effects of network topology such as signaling cascades, feedback loops and feed-forward loops on the propagation of chemical noise (17-22). These protein interaction networks often consist of multisite reversible covalent modifications of target proteins such as phosphorylation, acetylation, methylation, ubiquitylation. Such multisite modifications lead to change in catalytic activity, binding capability, transport or degradation of target proteins. Multisite phosphorylation is one such reversible covalent modification where a target protein is phosphorylated multiple times either by distributive or processive manner (23). When the same enzyme causes multiple phosphorylation in a distributive manner, it generates ultrasensitive signal response – a requirement for generating multistability and oscillations in signaling networks (23-25). Many crucial events in eukaryotic cell cycle are regulated by multisite phosphorylations of proteins Whi5, Cdh1, Sic1, Cdc25 by cyclin dependent kinase. Therefore, multisite phosphorylation mechanism has been widely used in mathematical and computational modeling of bistability and oscillations in cell cycle (26-30) – a fundamental property of any living organism. Therefore it is highly relevant to study the characteristics of chemical noise in a multisite phosphorylation chain. In particular study on the dependence of steady state variability of phosphorylated species on the various rate parameters of phosphorylation chain or the equilibrium constant of phosphorylation-dephosphorylation reactions.

Further, stochastic simulation algorithm (SSA) has been an obvious choice for accurate estimation of noise in chemical reaction networks of various cellular events (31). However due to the disparity in time scales of chemical reactions, particularly reactions with faster times scales tend to slow down the simulation as these reactions ‘fire’ recurrently in a given time interval. As phosphorylation/dephosphorylation of proteins usually occurs in faster time scales as compared to synthesis or degradation of proteins in living cells (32), computational cost of SSA are usually quite high in networks with multisite phosphorylation reactions. In order to improve the efficiency of SSA various methods have been developed exploiting the firing rate or the timescale separation of chemical reactions. Methods such as tau leaping, slow-scale SSA, stochastic quasi steady state approximation on enzyme catalyzed reactions are useful in this context with their own advantages and disadvantages (33-43) .

Keeping in view of the important role of multisite phosphorylation, in this paper we investigated two aspects of stochasticity in multisite phosphorylation chain. In order to investigate the quantitative and qualitative nature of variability in phosphorylated species, we used van Kampen’s system size expansion method to quantify noise at steady state. We examined the dependence of variability on the equilibrium constant of the reactions and on the total number of phosphorylation sites on the species. On account of timescale separation we applied quasi steady state approximation on the phospho chain and proposed a method to improve the computational efficiency of SSA. We applied the method on signal transduction cascade and positive feedback loop to validate the quantitative accuracy of the method.

## Methods

We studied noise propagation in ordered distributive multisite phosphorylation where each enzyme-substrate encounter leads to single phosphorylation or dephosphorylation of target protein and enzymatic events happen in a specific order(26). We present a three-component reversible distributive multisite phosphorylation scheme (Figure 1a). We considered that a single enzyme catalyzes all the phosphorylation events and similarly another enzyme catalyzes all the dephosphorylation events. In addition we also assumed that the enzymes are catalyze the forward and backward reactions are in much excess concentrations compared to their substrates such that the reactions can be represented by pseudo-first order mass action rate laws. The chemical master equation that three component reaction follows can be represented as:

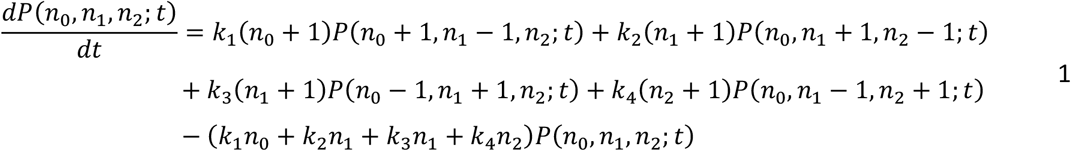

*P*(*n*_0_, *n*_1_, *n*_2_; *t*) represents the joint probability that the at time *t* there will be *n*_0_, *n*_1_ and *n*_2_ number of molecules for the species X_0_, X_1_ and X_2_ respectively. The total number of molecules of X, *n_T_* (= *n*_0_ + *n*_1_ + *n*_2_), is fixed and holds a mass conservation. Due to the mass conservation, the chemical master equation can also be written as a function of any two variables of the phospho chain (44).

In order to calculate the steady state fluctuations in phospho species we used linear noise approximation on master equation also known as van Kampen’s system size expansion or Ω expansion of master equation (45). Identifying Ω as the volume of the reaction chamber of the homogeneous reaction mixture, the main ansatz in the system size expansion is that the size of the fluctuations from the macroscopic average varies inversely with √Ω. Systematic expansion of master equation to the first order in 1/√Ω leads to macroscopic average equation. Whereas the expansion to second order in 1/√Ω generates a linear Fokker-Planck equation of the random variable. Since in the process of generating Fokker-Planck equation, higher order terms above 1/Ω^0^ are excluded it is called linear noise approximation. This method has been used extensively to calculate gene expression noise and noise in biochemical reaction networks (44, 46, 47).

**Figure 1:**
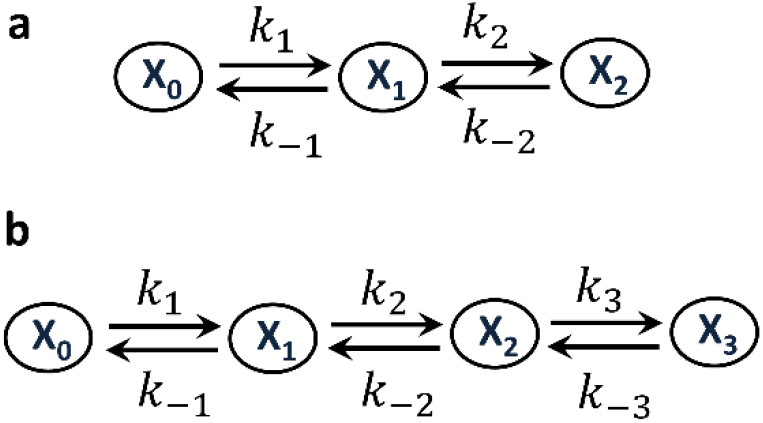
Schematic representation of three (a) and four component (b) phosphorylation chains. The rate constants of individual reactions are mentioned on top of arrows indicating chemical reactions. The subscript in the name of chemical species represents the number of phosphorylation.

At the steady state the drift and diffusion matrices in the Fokker-Plank equation are connected by a fluctuation-dissipation like relation given by

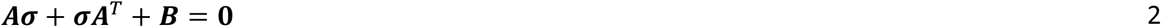

where ***A, B*** and ***σ*** are the drift or Jacobian matrix, diffusion matrix and covariance matrix respectively. The covariance matrix holds information about the variance and covariance of all the molecular species in the network. The elements in the drift matrix ***A*** are given by

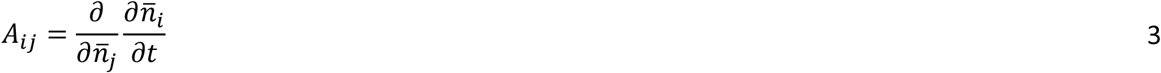

Where *n̅_i_* is the average number of molecules for the z’-th chemical species and can be obtained from the macroscopic rate equations (the derivative with respect to time on the right hand side). The elements in the diffusion matrix ***B*** are given by

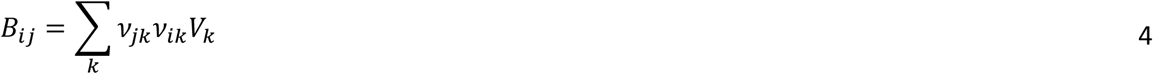

Where *v_ik_* and *V_k_* are the stoichiometric coefficient of the *i*-th species in the *k*-th reaction and the rate of the *k*-th reaction respectively. For the three component reaction scheme the drift and diffusion matrices are given by

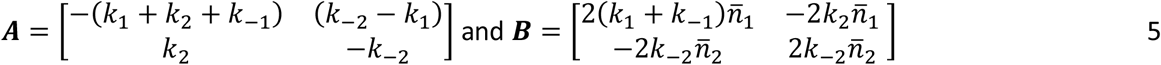

Using the above expressions of ***A*** and ***B*** in the fluctuation-dissipation relation (Eq.2) and applying the symmetry in covariance (*σ_ij_* = *σ_ji_*) we obtained a matrix equation for the steady state variances of stochastic variables as

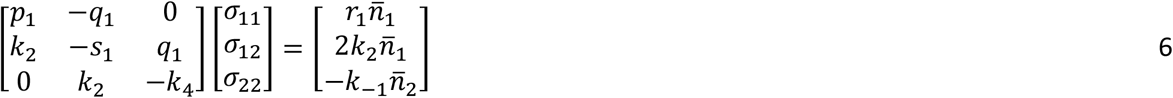

Where *p*_1_ = *k*_1_ + *k*_2_ + *k*_−1_, *q*_1_ = *k*_−2_ – *k*_1_, *r*_1_ = *k*_1_ + *k*_−1_ and *s*_1_ = *k*_1_ + *k*_2_ + *k*_−1_ + *k*_−2_. Note that as we have applied the mass conservation relation, the Eq.6 does not contain the variability term for the unphosphorylated species X_0_. Solution of the above system of linear equations lead to the variances of species X_1_ and X_2_ as

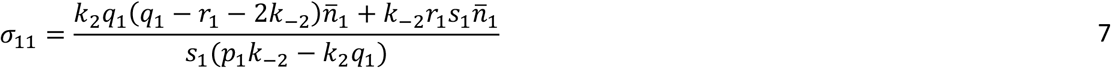

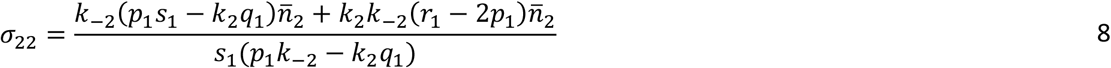

In the special case where all the forward and backward rate constants are equal i.e. *k*_1_ = *k*_2_ = *k_f_* and *k*_−1_ = *k*_−2_ = *k_b_*, the steady state variances can be represented as a function of equilibrium constant (*K* = *k_f_*/*k_b_*) and they take simple form as

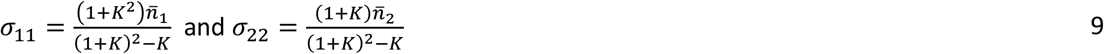

We proceed further to calculate steady state variances in a four-component equilibrium (Figure 1b). We followed the similar procedure as described for the three-component system. The Jacobian matrix is given by

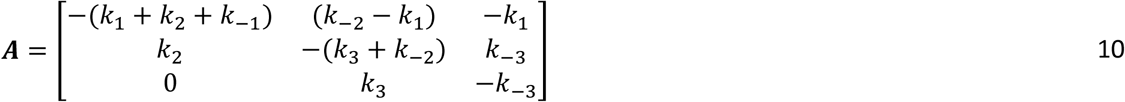

and the diffusion matrix is given by

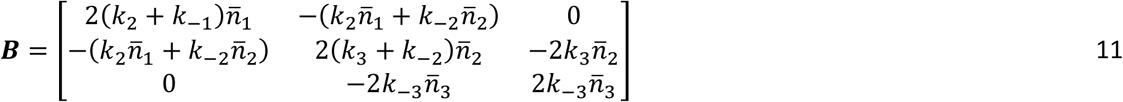

Making use of the fluctuation-dissipation relation (Eq. 2) we again obtained a matrix equation for the covariance as

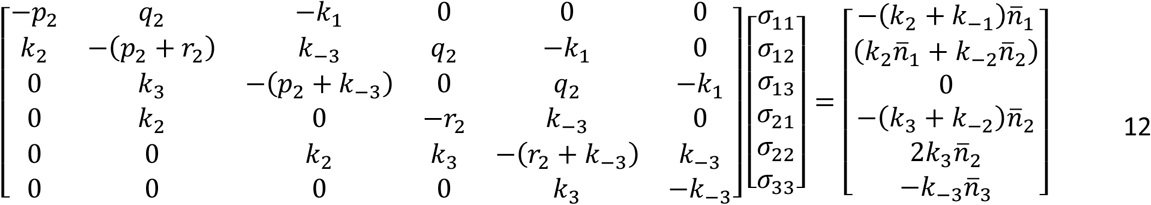

Solution of these linear equations provide the steady state variance of phospho species X_1_-X_3_. We have analytically solved these simultaneous linear equations to get the steady state variances. However due to complex nature of the solutions we prefer not to present them for the general case. However, in the special case where all the forward and backward rate constants are equal (*k*_1_ = *k*_2_ = *k*_3_ = *k_f_* and *k*_−_ = *k*_−2_ = *k*_−3_ = *k_b_*), the variances of X_1_, X_2_ and X_3_ can be expressed as a function of *K* as

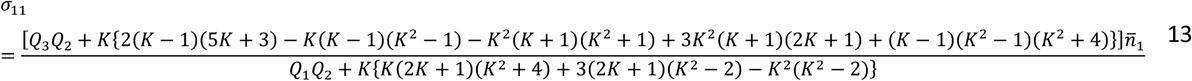

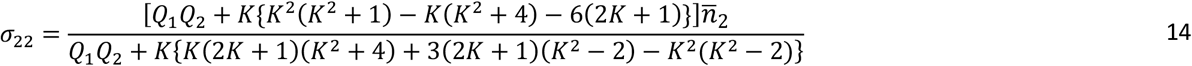

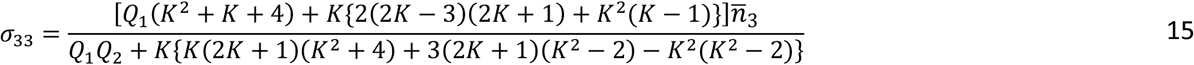

where *Q*_1_ = 2(*K* + 1)(2*K* + 1) + 3*K*^2^, *Q*_2_ = *K*^3^ + *K*^2^ + *K* + 4 and *Q*_3_ = (*K* + 1)(3*K* + 2) + 2*K*(*K*–1).

Although we have calculated only the variances using Eq. 12 one can also calculate covariances. Further the macroscopic averages can be obtained from the deterministic dynamical equations. The general expression of the average for *i*-th phospho state is given by (26)

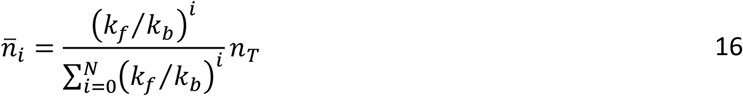

*N* is the total number of phosphorylation site in the chemical species.

## Results

In Figure 2 we present the dependence of steady state variance of phosphorylated species with the variation of rate constant of phosphorylation (left panel), dephosphorylation (middle) and equilibrium constant of phosphorylation-dephosphorylation reactions (right panel) for three (top row) and four (bottom row) component phospho chains. In case of three- and four-component chain, we used Eq. 9 and Eq. 13-15, respectively to get the equilibrium constant dependence on variance. To obtain the individual rate constant dependence for three-component chain we used Eq. 7-8 and for four-component chain we solved the matrix equation (Eq. 12) using Matlab. The variance increases sharply with increase of equilibrium constant and passing through a maximum it decreases with a long tail. The value of *K* at which the variance is maximum increases with the number of phosphorylation. Further comparisons with the numerical simulations of the chain using SSA show complete agreement of analytical and numerical calculations.

**Figure 2:**
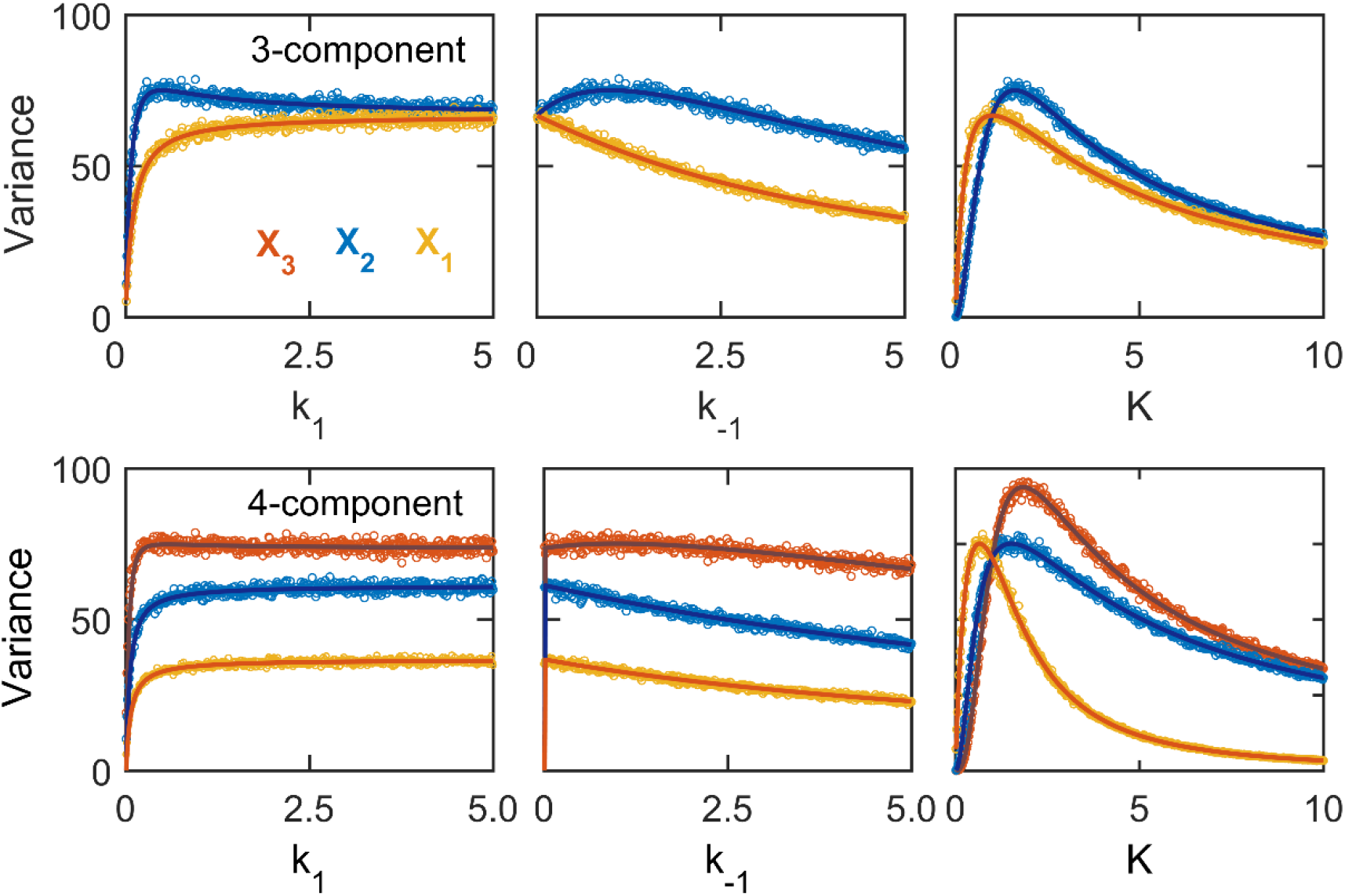
Dependence of steady state variance on the rate constants of chemical equilibria in phosphorylation chain. Solid lines and open circles represent analytical and numerical results respectively for three-(upper row) and four-component (bottom row) chains. Numerical results were from stochastic simulations of chain using SSA. Other numerical parameters were: for *k*_1_ variation *k*_2_ = *k*_3_ = 1 and *k*_−1_ = *k*_−2_ = *k*_−3_ = 0.5; for *k*_−1_ variation *k*_1_ = *k*_2_ = *k*_3_ = 1 and *k*_−2_ = *k*_−3_ = 0.5 and *n_t_* = 300.

Next we extended linear noise approximation calculation to five- and six-component phospho chains. Following the similar methodology as described for the previous cases we generated matrix equations for the covariance vector at the steady state. Applying the symmetry condition *σ_ij_* = *σ_ji_* the sizes of ***σ*** vectors for five- and six-component chains were 10 and 15, respectively. Therefore the resulting matrix equations were quite big and hard to solve analytically (see Supplementary text). To get the values of *σ_ii_* we solved these matrix equations using Matlab. We present the comparisons of analytical and numerical results of steady state variance in Figure 3. We further calculated coefficient of variation (*CV* = √*σ_ii_* / *n̅_i_*) for all the phospho species and compared with numerical results. CV provides information about the ‘noisiness’ exhibited by the chemical species in a network. Our calculations indicate that both for five- and six-component chains, below a critical value of equilibrium constant (*K* = 1) the noise increases with the number of phosphorylations and above the critical value the trend is reverse. Similar behavior was also obtained for three- and four-component chains (not shown). Therefore the crossover nature of noise with the variation of *K* is independent of the total number of phospho species in the chain. The crossover was due to the shift in distribution of abundance of phospho species with variation of equilibrium constant (26).

**Figure 3:**
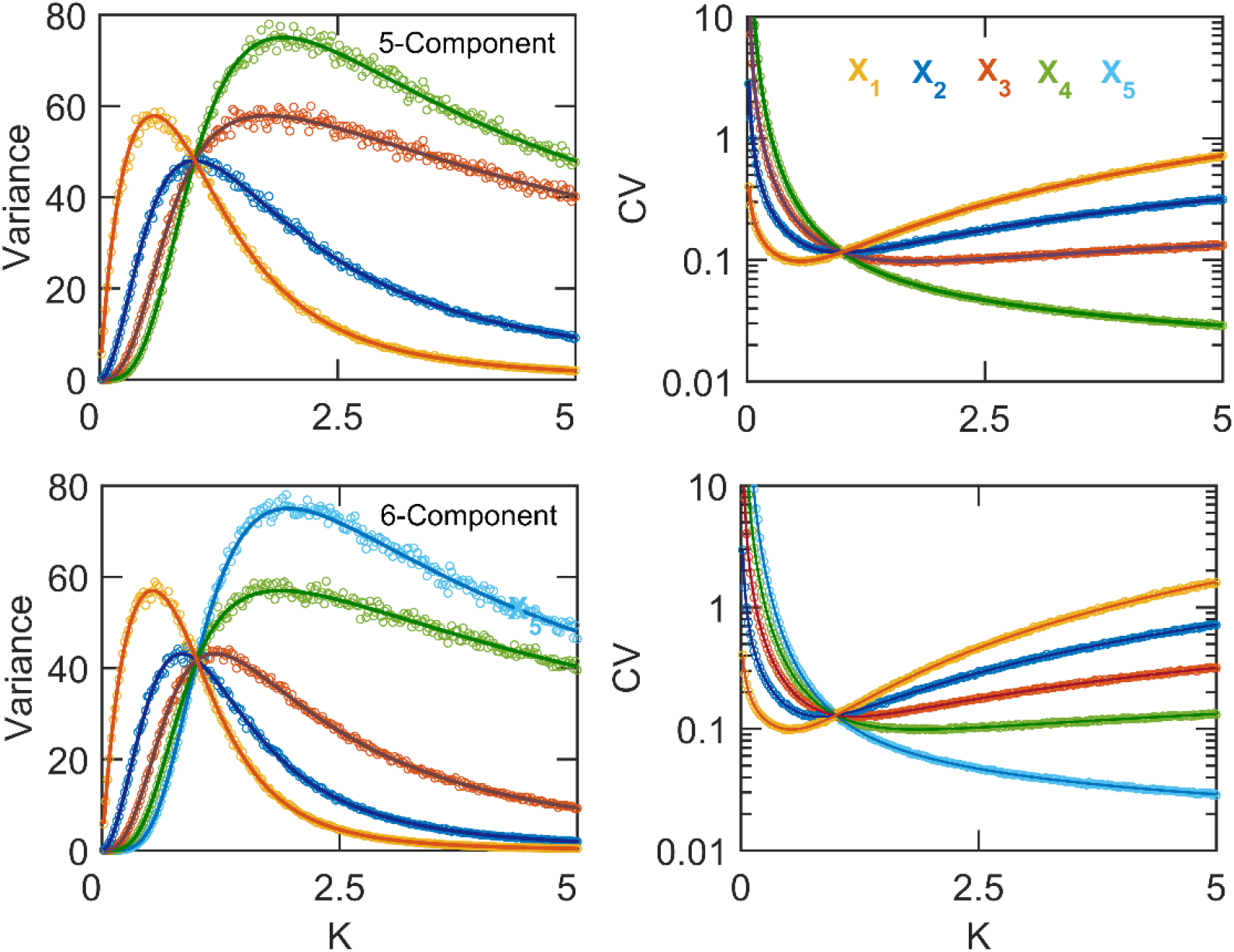
Comparison of steady state variance and CV obtained from analytical calculations (solid line) and numerical simulations (circle) for five-(upper row) and six-component (bottom row) phosphorylation chains. The total population (*n_T_*) of X was 300 in both the cases.

Many proteins in living cells are phosphorylated multiple times to generate threshold and ultrasensitive signal response in protein activity (48-50). Furthermore in a chain increased number of phospho states are known to cause sharper signal response curve with higher Hill coefficient values (23, 26). Keeping in mind this role of multi-phosphorylation, next we estimated the effect of increased phospho states on the noise of chemical species in a chain. In this regard we compared the analytically calculated CV of a given phospho-state (X_i_) from chains with different total number of phosphorylation sites or in other words the ‘length’ of the chain (Figure 4) keeping *n_T_* fixed. We found that for every phospho species the variability increased with increase of total number of phosphorylation sites. Therefore although higher number of phospho states are known to increase the ultrasensitive nature of signal response curve however our calculations indicated that it may lead to increase of variability in response. As we kept the total population (*n_T_*) same for all chains, with the increase of the number of states the average abundance of every phospho state decreased resulting in increased noise. Therefore a chain with more number of states may be nosier than a chain with less number of states. To determine the overall variability of the phospho states, we next calculated the total variance of all the phospho species by summing up their variances and found that the total variance increased with increase in the number of components in the chain (Figure 5).

**Figure 4:**
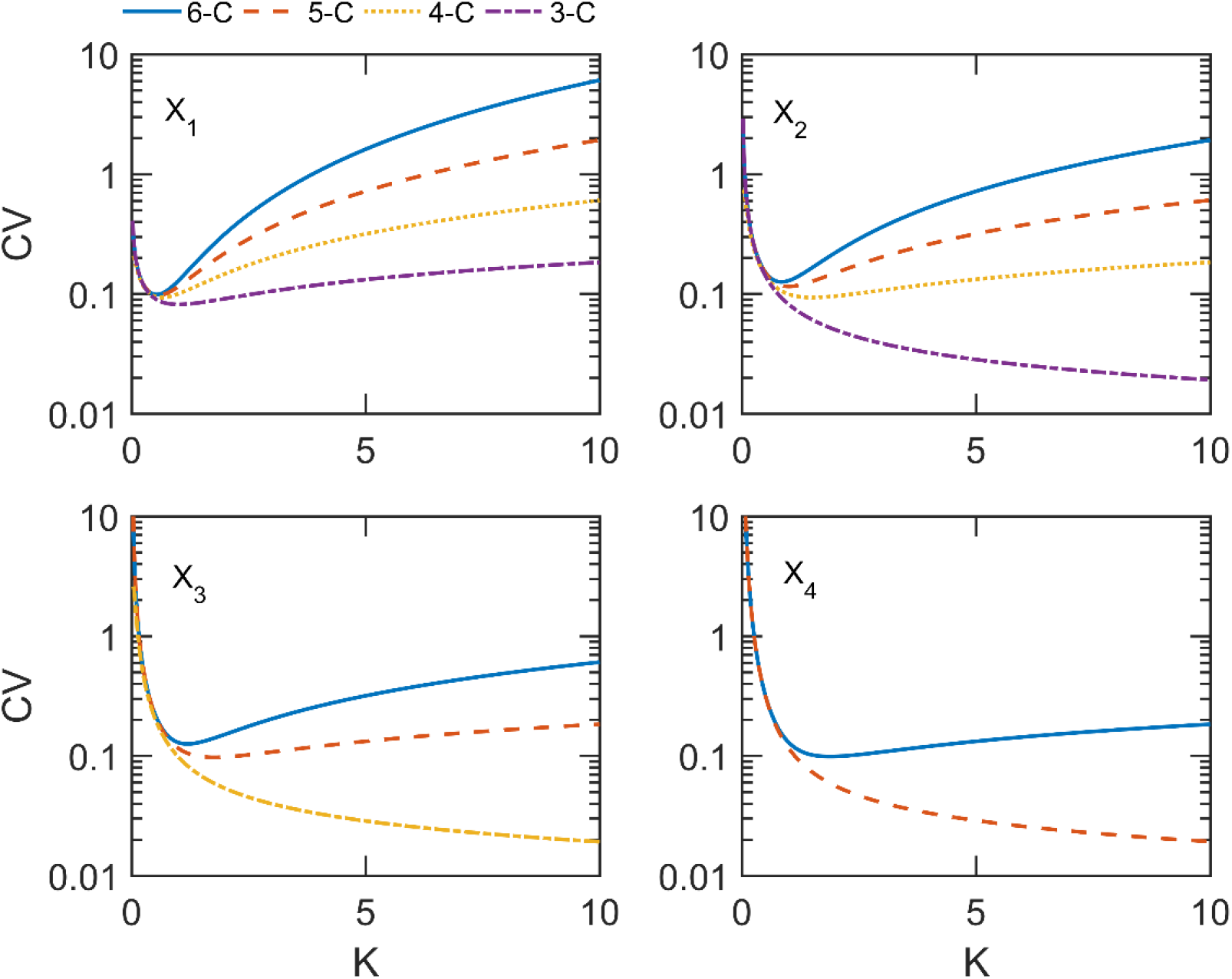
Comparison of noise in phosphorylated species from chains of different total number of phosphorylation states with *n_T_* = 300 for every chain. Different colors or line styles represent phospho chains with 3, 4, 5 and 6 number of components. The phospho states are indicated inside each plot.

**Figure 5:**
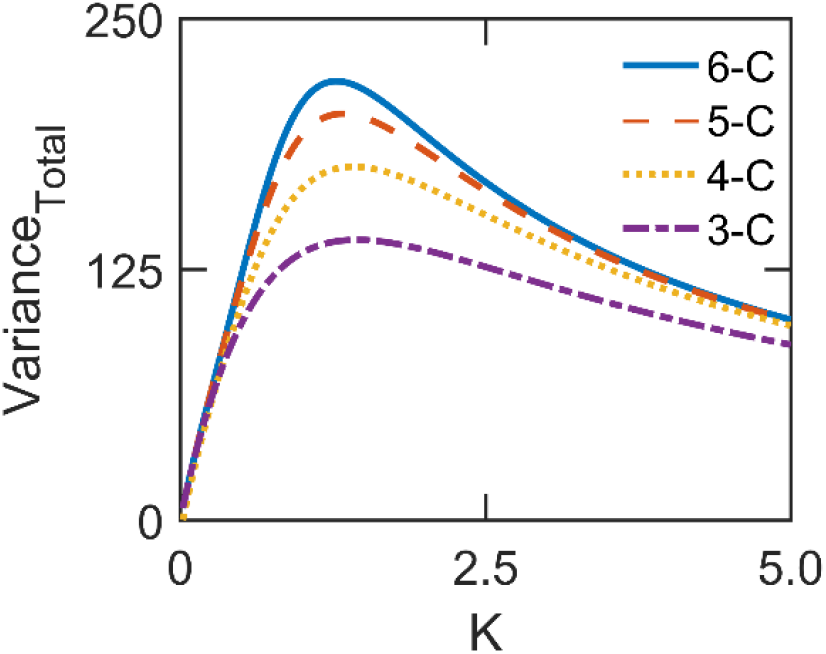
Variation of the total variance (=∑ *σ_ii_*) with the equilibrium constant. Each color or line style represents chain with different total number of phosphorylation sites.

### Application: quasi steady state approximation

Population abundance heterogeneity and disparity in timescales in coupled chemical reactions often pose a significant challenge towards computational efficiency of SSA (51). For a coupled chemical reaction network involving multi-phosphorylation chain, we applied quasi steady state approximation on the phospho chain under the condition of time scale separation between the reactions in the chain and the rest of the network. Specifically when the reactions in phospho chain occur in much faster timescale as compared to other reactions in the network, it allows the conditional probability distribution of fast variables to reach a quasi steady state in the timescale of other reactions. This is stochastic analogue of deterministic quasi steady state approximation (52). Under this condition the conditional probability distribution reaches a stochastic quasi steady state (36, 39, 40)

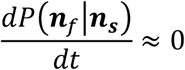

Where ***n**_f_* and ***n**_s_* are the population of fast and slow variables, respectively. The QSSA assumption on the probability leads to reduction in dimension of the reacting system (40, 53). Thus the reactions evolving in slower time scales are simulated using SSA where the propensities of slow reactions depend parametrically on the steady state average of the fast variables obtained from the steady state solution of their macroscopic dynamical equations. However as the values of the fast variables are taken from the deterministic equations, they lack the effect of intrinsic noise of chemical reactions. Therefore in order to include the effect of intrinsic noise in the fast variables, at each instant to the deterministic average of fast variables we added Gaussian white noise terms. The added noise was sampled from a Gaussian distribution with zero mean and the variance was taken from the analytical solution provided by the linear noise approximation of the chain. For example in a network with four-component phospho chain, for the species X_3_ we added a zero mean Gaussian noise having variance given by *σ*_33_ in Eq. 15. The result of linear noise approximation is a Fokker-Planck equation having a solution of Gaussian form for the probability distribution of random variables. Therefore adding a Gaussian noise with the variance given by analytical solution of fluctuation-dissipation relation is justified. The steps of modified QSSA were as follows:

1. Initialized of all chemical species.
2. Calculated of steady state deterministic average (*n̅_i_*) and variance (*σ_ii_*) of phospho species using the analytical expressions given by linear noise approximation.
3. Added Gaussian noise to the deterministic average: 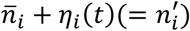 where *η*(*t*) is a Gaussian white noise having statistical properties: <*η_i_*(*t*)> = 0 and <*η_i_*(*t*)*η_i_*(*t*’) > = √*σ_ii_δ*(*t* – *t*’).
4. Calculated the propensities of all chemical reactions in the reduced system i.e. excluding the reactions of the chain; calculated next reaction time interval following SSA method and updated the time.
5. Determined the next reaction and updated the species abundance as per the stoichiometry of the chemical reaction occurred.
6. Repeated the process from step 2.

Using this recipe we applied our proposed method in linear cascade signal transduction network and positive feedback loop – two commonly found reaction motifs in living cells.

We first applied QSSA method to a signal transduction cascade network where information flows only in one direction without any feedback loop. The network diagram is given in Figure 6a. A four-component phospho chain was embedded in the network where the species Y acted as the enzyme catalyzing phosphorylation reactions in the chain and only the terminal phosphorylated species, X_3_, was assumed to catalyze synthesis of the species Z. The full network consisted of 10 chemical reactions and upon QSSA approximation the reduced system consisted only 4 chemical reactions associated with the synthesis and degradation of Y and Z (Table 1). At each time for a given value of Y, average value of X_3_ was calculated using the relation given by Eq. 16. A Gaussian random number with zero mean was added to the average of X_3_ where the variance of the random noise was given by the Eq. 15. Further, to investigate the effect of different total number of X on the variability, the zeroth and second order rate constants were multiplied and divided by *n_T_*, respectively leading to the appropriate scaling of the system. Note that in the average and variance equation (Eq. 15 and Eq. 16) *k_f_* was replaced by *k_f_n_Y_*/*n_T_* due to the second order nature of phosphorylation reaction. In Figure 6b we present a comparison between steady state CV of X_3_ and Y calculated using full SSA and QSSA methods for different values of synthesis rate of Y (*k_s_*). We further extended the comparison for different values of the equilibrium constant of the chain (Figure 6b-d) by varying *k_b_* keeping *k_f_* constant. We found that that in all cases the QSSA accurately recaptures the noise in all the regulators in the network. The accuracy of QSSA method was limited if the Gaussian noise was not added to the average of X_3_ (Supplementary Figure S1). Further in order to quantitatively estimate the accuracy of the QSSA method, we estimated a deviation error by calculating the sum-squared-deviation (=∑*_i_*(*x_i,QSSA_* – *x_i,SSA_*)^2^). The deviation error in CV was quite less (<10^-3^) both for X_3_ and Y (Figure 6e).

**Figure 6:**
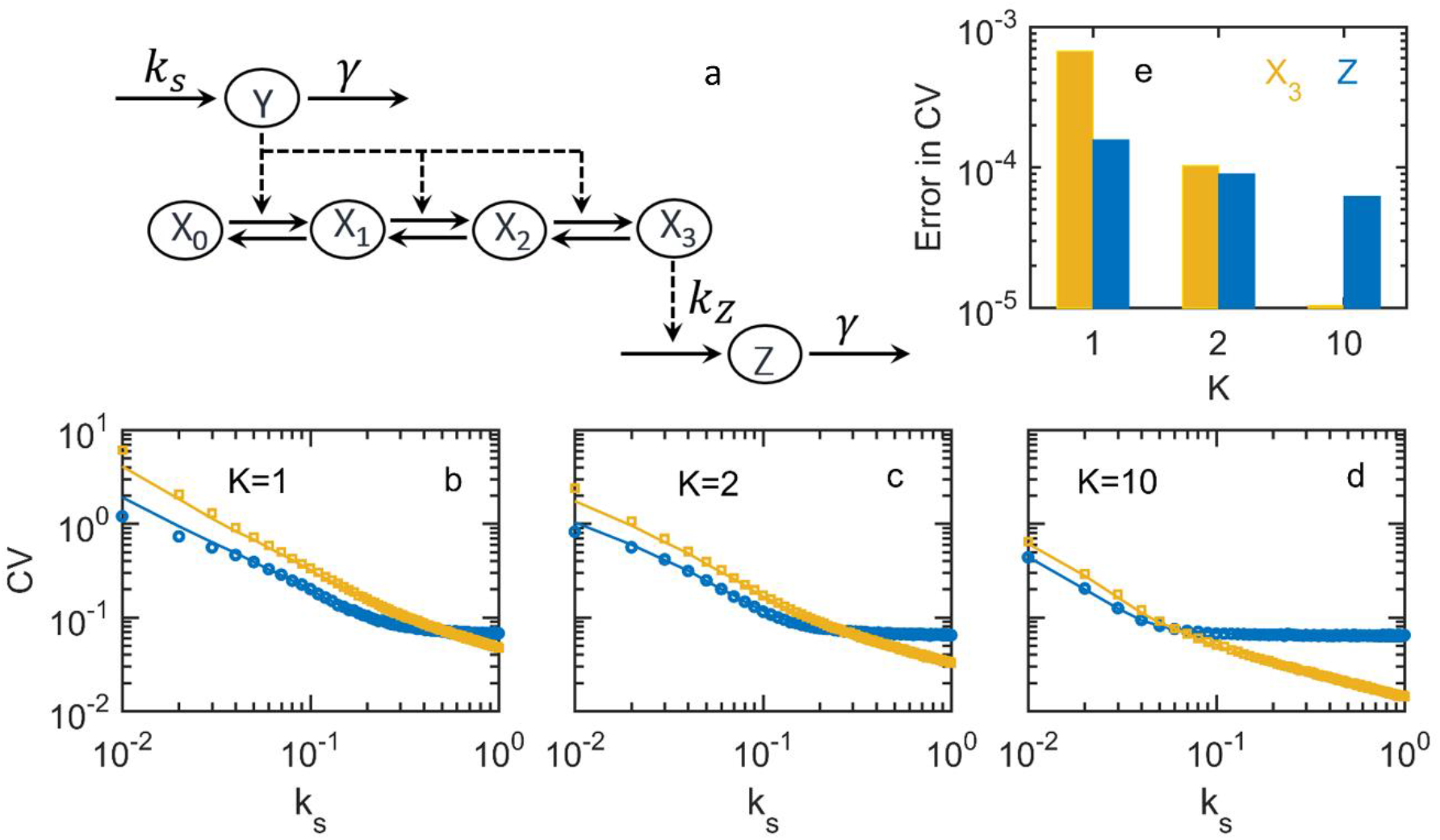
a) Schematic diagram of signal transduction cascade model where Y catalyzes phosphorylation of X and X_3_ catalyzes production of Z. Chemical reaction and catalytic activity to a reaction are represented by solid and dashed arrows, respectively. (b-d) Comparison of CV from full SSA (solid line) and QSSA (circle or square) simulations for X_3_ (circle) and Y (square) for indicated values of *K*. (e) Error in CV, defined as sum-squared-deviation QSSA from full SSA, for different values of *K*. The parameter values used were: *γ* = 0.1, *k_z_* = 0.5, *k_f_* = 10 and *n_T_* = 50. To vary the equilibrium constant different values of *k_b_*(= 10, 5,1) were chosen.

**Table 1:**
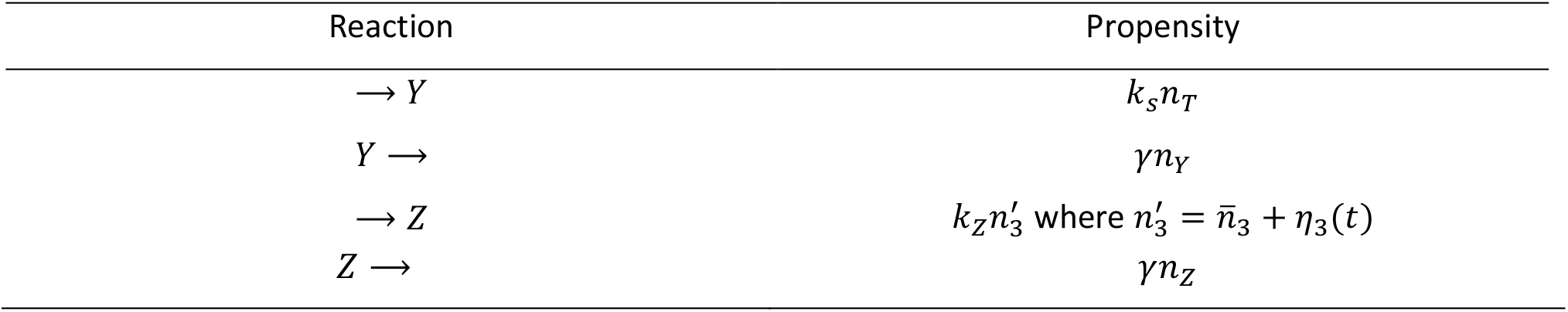
List of chemical reactions in the reduced system of signal transduction cascade network. *η_3_*(*t*) is a Gaussian random number following <*η_3_*(*t*)> = 0 and < *η_3_*(*t*)*η_3_*(*t’*) > = √*σ_33_δ*(*t* – *t*’).

We next applied the QSSA method to a positive feedback loop network that generate bistable signal response. In living cells bistability is quite a common feature of chemical reaction network with positive feedback loop. All-or-none type of response generated by a bistable switch helps cell in making various decision making processes in proliferation, death and differentiation (54). Previously multisite phosphorylation in a positive feedback loop has been predicted to generate bistability even with mass action reaction rates in the chain (26). Here, as in previous network Y acted as kinase to X and the terminally phosphorylated X_3_, assumed to be active from of X, catalyzed synthesis of Y thereby creating a positive feedback loop between X and Y (Figure 7a). On application of QSSA on the chain, the reduced system consisted only two reactions associated with the synthesis and degradation of Y with propensities 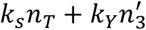 and *γn_Y_*, respectively. We made a comparison of steady state population of Y generated from full SSA and QSSA of the model while the bifurcation parameter, the unregulated synthesis rate constant of Y (*k_s_*), was varied (Figure 7b-c). Here the steady state distribution of Y is the stochastic analogue of the deterministic one-parameter bifurcation diagram that was overlaid on top of the population distributions. We made the comparison initializing the simulations at the lower (Figure 7b) and the upper steady states (Figure 7c) of the deterministic bifurcation diagram and in both cases results from QSSA simulations were almost identical to the full SSA. A full comparison of steady state distribution of Y across the bifurcation diagram can be found in the Supplementary Figure S2. The distributions obtained from QSSA simulations were quite similar to the distributions from SSA simulations. For quantitative comparison of noise, next we calculated average and CV of Y for three different values of *n_T_* that scales the abundances of all the species without shifting the bifurcation diagram in x-axis (Figure 8). Again we found that QSSA method was able to capture noise accurately in all three cases. The comparison at different values of *n_T_* highlights the important fact that even with low population abundance (*n_T_* = 50), the QSSA method was able to capture the noise accurately. Further the accuracy of QSSA does not depend on the initial conditions used in the simulations of bistable switch.

**Figure 7:**
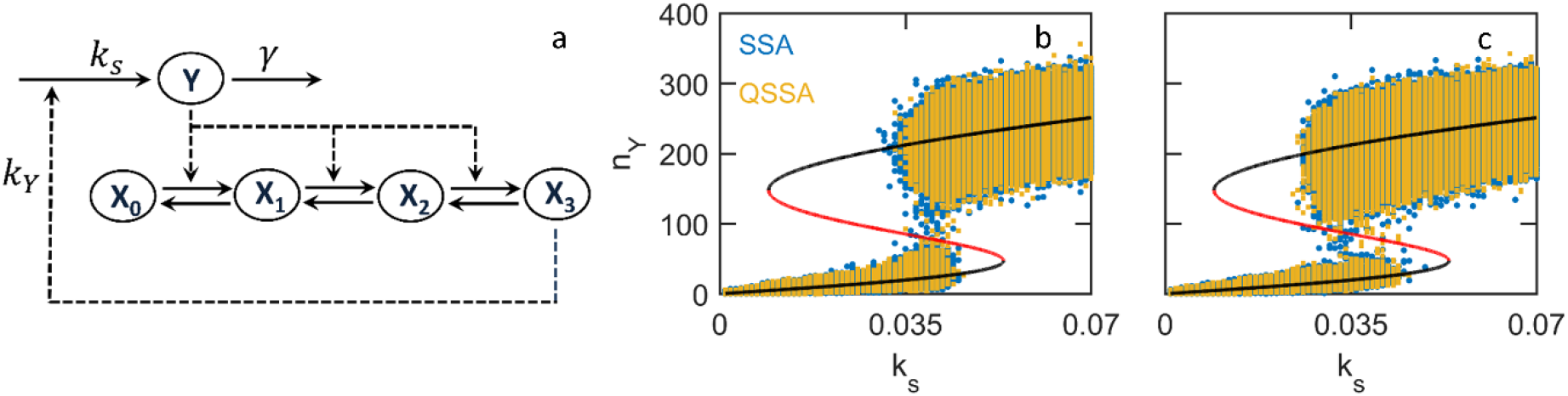
Schematic diagram of a positive feedback loop involving 4-component multi-phosphorylation chain (a). Comparison of steady state populations of Y from full SSA and QSSA simulations initialized from lower (b) and upper (c) steady states of deterministic bifurcation diagram. The sloid line represents one-parameter bifurcation diagram of the model. The stable and unstable steady states are represented by black and red lines, respectively. The parameter values used were: *γ* = 0.1, *k_Y_* = 0.7, *k_f_* = 1, *k_b_* = 2 and *n_T_* = 50.

So far we assumed that the terminally phosphorylated species X_3_ was the only active form X. However in many protein regulatory networks two or more phospho species of same protein may act as active form of the protein. Therefore in order to investigate the effect of other phospho species on the accuracy of QSSA, next we assumed that both X_3_ and X_2_ are capable of catalyzing the production of Y. Now in this case the propensity of production of Y was given by 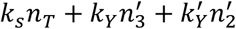. A comparison of full SSA and QSSA showed that in this case also QSSA accurately recaptures the noise in Y (Figure 9). Note that the region of bistability in this case was a bit smaller (Supplementary Figure S3) as compared to the previous case due to decreased ultrasensitivity in the chain (26). Further to investigate the effect of time scale separation, we performed full SSA simulation of the model with reduced values of both *k_f_* and *k_b_* keeping the equilibrium constant of the chain same. We found that by slowing down the reactions in the chain, the noise in Y changes significantly both for simulations initialized at the upper and lower (Supplementary Figure S4) steady states. Therefore the accuracy of QSSA would critically depend on the timescale separation which requires faster *k_f_* and *k_b_* in the chain of phosphorylation-dephosphorylation reactions.

**Figure 8:**
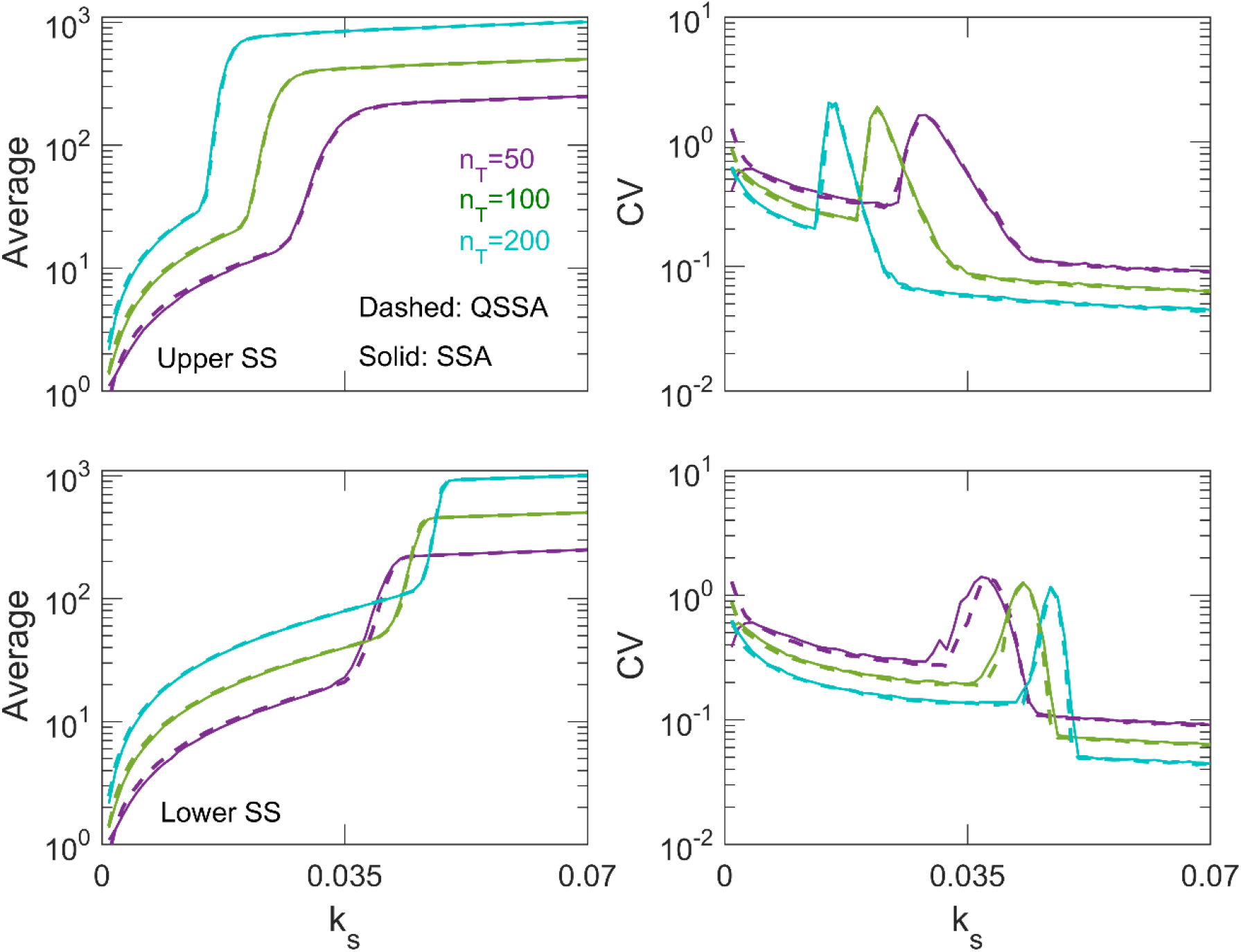
Comparison of steady state average and CV of Y for the indicated values of *n_T_*. Upper and lower panels are results from simulations initialized at the upper and lower steady states of the bifurcation diagram, respectively. The parameter values are same as Figure 7.

**Figure 9:**
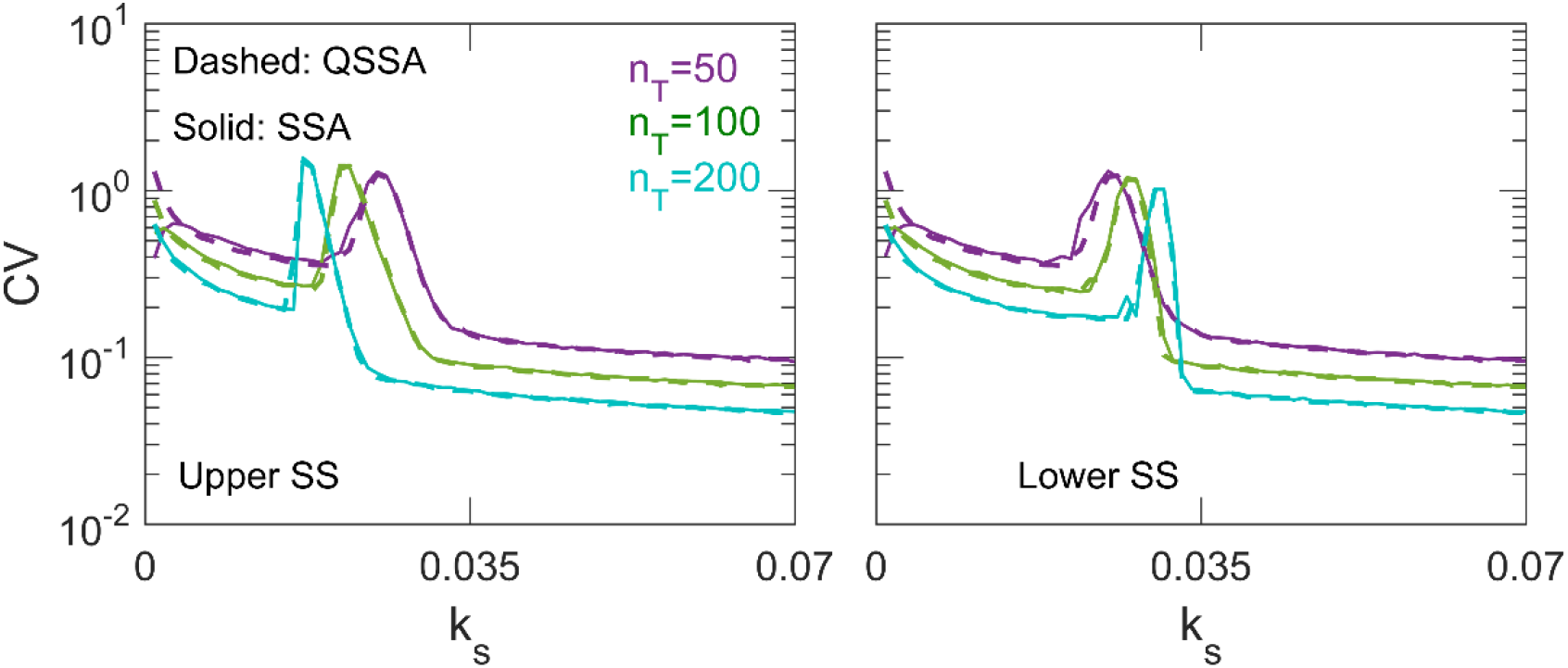
Upper row: Comparison of CV from full SSA and QSSA for the positive feedback network where both X_3_ and X_2_ catalyzes production of Y. Parameter values were: *k_Y_* = 0.5, 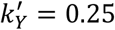 and others were same as Figure 7.

## Conclusion

Chemical reactions in living cells experience unavoidable chemical noise resulting in cell-to-cell variability of cellular functions or characteristics. The origin of cell-to-cell variability was found to be the gene expression noise whose propagation critically depends on the topology of network. Multisite phosphorylation of protein is often responsible for activation, inactivation, recognition, binding, degradation of target proteins. Further it is also responsible for ultrasensitive signal response in signaling network. Therefore understanding of chemical noise propagation in a chain of multisite phosphorylation reactions is important due to its vast role in cellular signaling. In this paper we analytically investigated chemical noise propagation in a multisite phosphorylation chain using van Kampen’s system size expansion method. We calculated steady state variance of phosphorylated species while we varied the equilibrium constant of phosphorylation-dephosphorylation reaction and the total number of phosphorylated states in the chain or the chain length. The variability or noise in phospho species, measured by the coefficient of variation, showed a crossover at a critical value of equilibrium constant (*K* = 1). Below the critical value the variability increases with increasing number of phosphorylations whereas above the critical value the variability decreases with increasing number of phosphorylations. This behavior is consistent for chains of different lengths. The noise at a given phosphorylated state depends on the total number of phosphorylated states in the chain - noise increases with increasing number of states in the chain. Further we found that the total variance of all the phosphorylated species increases with increasing number of states in the chain. Previous studies have shown that more phosphorylated states (or ‘longer’ chain) lead to increased ultrasensitive nature of signal response curve(25, 26), however our calculations indicate that it may result in increase of variability in the response.

Next we made use of the steady state variability given by linear noise approximation to propose a recipe for improving computational efficiency of stochastic simulation algorithm applying a quasi steady state approximation on the phosphorylation chain assuming a timescale separation between the phospho chain and the rest of the network. The key ingredient in the method is that a zero-mean Gaussian white noise was added to the steady state average of phospho species in order to include the effect of inherent stochasticity of phosphorylation-dephosphorylation chain. The variance of Gaussian noise was given by the analytical solution of fluctuation-dissipation relation from linear noise approximation. We applied the recipe to a signaling cascade and to a positive feedback loop network motifs. We found that under the condition of timescale separation our recipe estimated the average and the noise accurately in these systems. As our analytical method correctly estimated the noise in the phospho species, the calculated variability in the downstream signaling was accurate in QSSA method. As application of QSSA lead to dimensional reduction of the system, the number of reactions fired in a given time interval reduced significantly. For example in the positive feedback loop network with a total of 50 molecules in the chain (*n_T_* = 50) the average number of reactions fired were ~3.6×10^5^ and ~2.2×10^6^ for QSSA and full SSA methods respectively till a simulation time of 10000 arbitrary unit in the bistable region (*k_s_* = 0.035). In the same situation with 100 total molecules in the chain, the number of fired reactions were ~8.3×10^5^ and ~4.6×10^6^ for QSSA and full SSA methods, respectively. Due to the less number of reactions the QSSA method was computationally inexpensive as compared to the full SSA method. In the mentioned cases QSSA method took about 3 and 7 minutes for 50 and 100 molecules in the chain, respectively. Whereas full SSA method spent about 9 and 17 minutes, respectively. Therefore the proposed recipe lowers the computational cost significantly without compromising the accuracy of calculation. Particularly the method will be quite useful for networks where the simulation spends most of the time in simulating fast reactions.

## Acknowledgments

The work was supported by funding from the Science and Engineering Research Board, Department of Science and Technology (India), grant no. EMR/2015/001899, to DB. SD acknowledges fellowship from INSPIRE program of Department of Science and Technology, India.

